# Generation of a novel Nkx6-1 Venus fusion reporter mouse line

**DOI:** 10.1101/2021.01.29.428800

**Authors:** Ingo Burtscher, Marta Tarquis-Medina, Ciro Salinno, Julia Beckenbauer, Mostafa Bakhti, Heiko Lickert

**Affiliations:** Institute of Diabetes and Regeneration Research, Helmholtz Zentrum München, 85764 Neuherberg, Germany; German Center for Diabetes Research (DZD), D-85764 Neuherberg, Germany; Technische Universität München, School of Medicine, 81675 München, Germany

**Keywords:** Nkx6-1, pancreas development, fluorescent reporter, endocrine lineage, secondary transition, β-cells, live-imaging

## Abstract

Nkx6-1 is a member of the Nkx family of homeodomain transcription factors (TF) that regulates motor neuron development, neuron specification and pancreatic endocrine and β-cell differentiation. To facilitate the isolation and tracking of Nkx6-1-expressing cells, we have generated a novel Nkx6-1 Venus Fusion (Nkx6-1-VF) reporter allele. The Nkx6-1-VF knock-in reporter is regulated by endogenous cis-regulatory elements of Nkx6-1 and the fluorescent protein fusion does not interfere with the TF function, as homozygous mice are viable and fertile. In addition, the nuclear localization of Nkx6-1-VF protein reflects the endogenous Nkx6-1 protein distribution. During embryonic pancreas development, the reporter protein marks the pancreatic ductal progenitors and the endocrine lineage, but is absent in the exocrine compartment. Moreover, the levels of Nkx6-1-VF reporter is upregulated upon β-cell differentiation during the major wave of endocrinogenesis. In the adult islets of Langerhans, the reporter protein is exclusively found in insulin-secreting β-cells. Importantly, the Venus reporter activities allows successful tracking of β-cells in live-cell imaging and their specific isolation by flow sorting. In summary, the generation of Nkx6-1-VF reporter line provides a unique tool to study the spatio-temporal expression pattern of this TF during organ development and isolate and track Nkx6-1-expressing cells such as pancreatic β-cells, but also neurons and motor neurons in health and disease.

## Introduction

The endoderm-derived pancreas comprises exocrine and endocrine compartments that contribute to nutrient digestion and regulate blood glucose homeostasis, respectively. The exocrine pancreas consists of ductal epithelium and acinar cells, whereas the endocrine pancreas resides in the islets of Langerhans and contains five distinct hormone-producing cell types (Pan & Wright, 2011a; Shih et al., 2013). Among these, insulin-producing β-cells are the most prominent pancreatic endocrine cell type that represents around 80% of the total endocrine population in the mouse adult islets (Brissova et al., 2005; Roscioni et al., 2016). In mice, pancreas organogenesis is initiated by the formation of a dorsal and a ventral pancreatic bud from the foregut endoderm into the surrounding mesenchyme at embryonic day 9 (E9.0). The plexus structure of the early buds consists of multipotent progenitor cells (MPC), which are characterized by the expression of several transcription factors (TFs) including Pdx1, Ptf1 and Nkx6-1. From E9.5 to E12.5, the pancreas undergoes the primary transition, in which progenitor cells massively proliferate and form a transiently stratified epithelium surrounding microlumen structures. During the secondary transition (E13.5-15.5), the pancreatic epithelium undergoes extensive remodeling to generate a continuous tubular network followed by enormous cell differentiation to produce ductal, acinar, and endocrine lineages. At E15.5, the mouse pancreas contains a ramified tubular epithelium, which consists of round tip domain containing acinar progenitors (Ptf1^+^) and a trunk domain (Sox9^+^). Within the trunk region bipotent progenitors possess ductal or endocrine fate potential (Aimeé Bastidas-Ponce et al., 2017; Pan & Wright, 2011b; Villasenor et al., 2010). Upon differentiation, the endocrine cells leave the ductal epithelium and assembled into clusters to form the islets of Langerhans (Kesavan et al., 2014).

The Nkx family of homeodomain factors includes several members, among which Nkx6-1 plays a key role during foregut patterning, pancreas organogenesis, and central and peripheral nerve system development. In the ventral neuronal progenitors, Nkx6-1 plays a role in progenitors specification by modulating neural response to the glycoprotein sonic hedgehog (Shh) signals (Li et al., 2016; Prakash et al., 2009). In the foregut, Nkx6-1 is expressed in the smooth muscle cells of esophageal and dorsal tracheal mesenchyme and its function is required for promoting smooth muscle development in the esophageal region (Cai et al., 2000). During pancreas development, Nkx6-1 plays a key role in pancreatic epithelium patterning. Reciprocal suppression of Nkx6-1 and Ptf1a leads to the formation of the trunk and tip domains, respectively (Schaffer et al., 2010; Zhou et al., 2007). Therefore, upon loss of Nkx6-1 acinar fate is increased (Schaffer et al., 2010) at the expense of endocrine cells (Nelson et al., 2007; Sander et al., 2000; Schisler et al., 2008; Tessem et al., 2014). As the differentiation of bipotent cells into β-cells proceeds, the levels of Nkx6-1 increases (Øster et al., 1998). Together with Pdx1, Nkx6-1 is required for β-cell specification by preventing the α-cell program (Liu et al., 2011; Schaffer et al., 2013). In adult mice, Nkx6-1 is exclusively expressed in β-cells (Jensen et al., 1996) to maintain their identity and proper insulin secretion (Schaffer et al., 2013; Taylor et al., 2013). Lack of Nkx6-1 in mature β-cells leads to rapid-onset of diabetes caused by defects in insulin biosynthesis and secretion without affecting cell survival (Taylor et al., 2013). Furthermore, decreased expression of NKX6-1 is associated with development of T2D in humans and rodents (Guo et al., 2013; Talchai et al., 2012). In addition, three genome-wide association studies (GWAS) have identified NKX6-1 variants associated with T2D (Spracklen et al., 2020; Suzuki et al., 2019; Yokoi et al., 2006), further suggesting the importance of this TF for human β-cell formation and function.

Here, we have generated a novel reporter mouse line, in which the bright fluorescent protein Venus is fused to the C-terminus of the endogenous Nkx6-1. The reporter mice are viable and fertile. The Nkx6-1-Venus fusion (Nkx6-1-VF) protein follows the spatio-temporal expression pattern of endogenous Nkx6-1 during pancreas development and in adult islets. Moreover, the expression of Venus enables one to track β-cells in live imaging and isolate them specifically by FACS. Thus, the Nkx6-1-VF mouse line provides a unique tool to study Nkx6-1 expression and function during organ development as well as β-cell function in health and disease.

## Results and discussion

### Generation of the Nkx6-1-VF Mouse Line

The Nkx6-1 Venus fusion mouse line was generated by Crisp/Cas9-mediated double strand breaks followed by homologous recombination. Using a targeting vector as template DNA directed repair resulted in the generation of a *Nkx6-1-VF* reporter gene (Nkx6-1-VF) under control of the endogenous *Nkx6-1* promoter (Fig. 1a). To do so, we designed a targeting vector by standard cloning techniques, removing the translational stop codon of the *Nkx6-1* gene in exon 3 and generating an in-frame fusion transcript with the Venus open-reading frame and a Flag tag. For selection purposes, a loxP-flanked phospho-glycerate kinase (PGK) promoter-driven neomycin (neo) resistance gene in the opposite orientation was inserted after the reporter gene. After removal of the neo selection marker, the *Nkx6-1-VF* mRNA transcript utilizes the endogenous untranslated region (UTR). The targeting vector, Cas9D10A expression vector and two gRNA vectors expressing gRNAs that bind shortly before and after the *Nkx6-1* stop codon were electroporated into F1 hybrid (129Sv/Bl6) IDG3.2 ES cells. Neomycin resistant clones were screened with 5’ and 3’ homology arm spanning PCR (Fig. 1b and 1c). Germline chimeras of the *Nkx6-1-VFneo* mouse line were generated from two independent ES cell clones by ES cell aggregation with CD1 morulae. The loxP-flanked neo selection cassette was deleted in the germline by Cre recombination mediated excision (Fig. 1d; Soriano, 1999) resulting in the Nkx6-1-VF mouse line. Heterozygous animals were intercrossed and genotyped for all possible alleles (Fig. 1e). No embryonic phenotype was observed and pubs homozygous for the Nkx6-1-VF allele were viable and appeared indistinguishable from their wild-type littermates. To confirm the Nkx6-1-VF reporter is properly synthesized and shows similar characteristics as the endogenous Nkx6-1 protein, we performed Western blot analysis using lysates from islets of Langerhans from WT, heterozygous and homozygous mice. The Nkx6-1 WT protein was detected as a double band at 50 and 53 kDa with the anti-Nkx6-1 antibody (Fig. 1f). This double band was also observed with the Nkx6-1-VF protein appearing as a double band at approximately 77 and 80 kDa using anti-Nkx6-1, anti-GFP and anti-FLAG antibodies (Fig. 1f-h). Both the Nkx6-1 and Nkx6-1-VF protein were synthesized in comparable ratios, as revealed by anti-Nkx6-1 antibody in lysates from heterozygous animals (Fig. 1f). As such, the Nkx6-1-VF reporter protein can be used to quantify the Nkx6-1 protein levels in a molar ratio, to analyze cell-fate decisions in time-lapse studies, purify cell populations by fluorescent activated cell sorting (FACS), but also to analyze microRNA effects on the UTR using the Venus reporter as a live sensor for protein translation.

**Figure 1.**
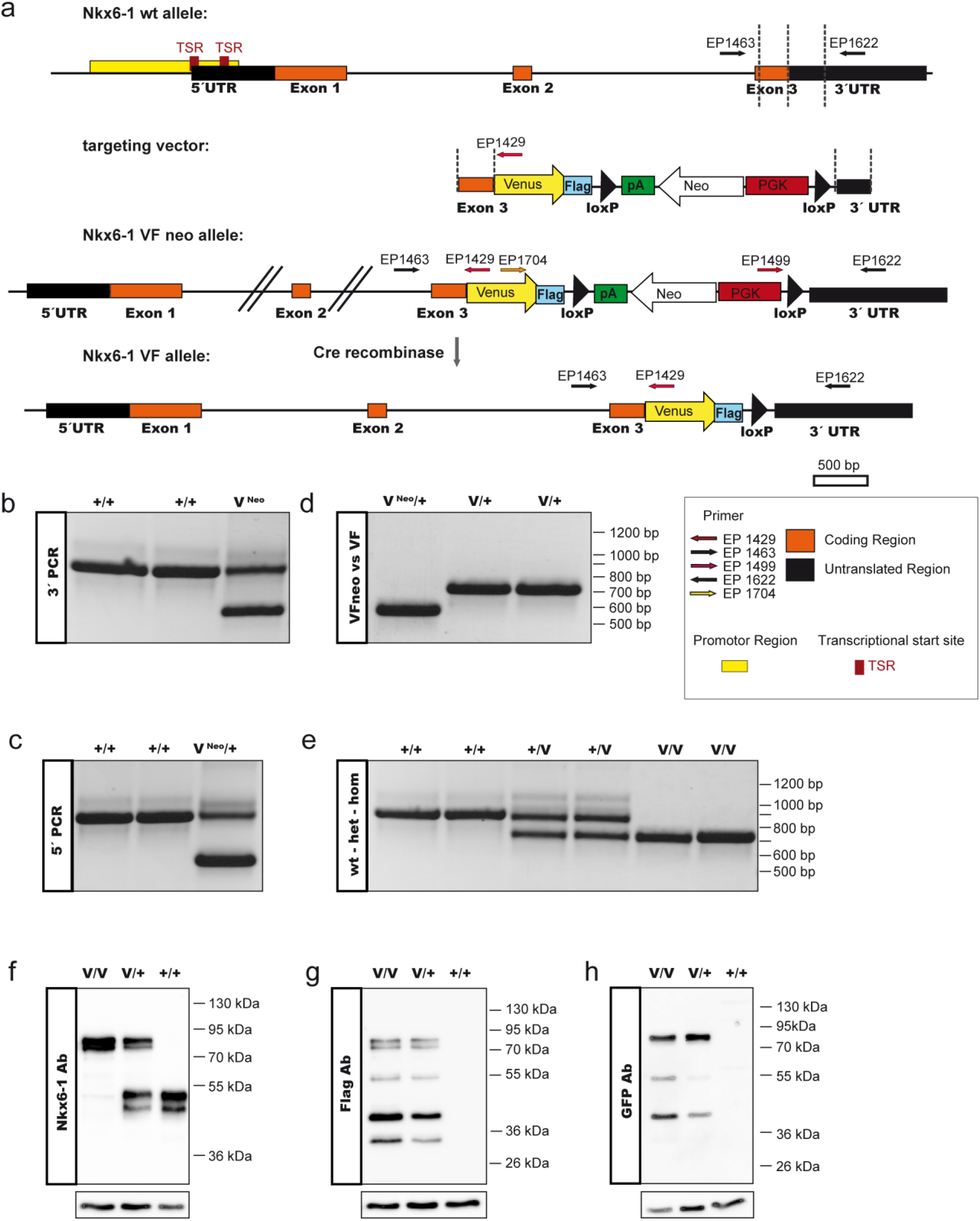
Generation of the Nkx6-1-VF allele. (**a**) Targeting strategy for the Nkx6-1-VF allele. A double strand break was introduced by two nicks of the D19A mutant Cas9 using two gRNAs (green arrows) binding before and after the stop codon of Nkx6-1. A targeting vector was used to repair the gap and fuse the coding region of the fluorescent report gene Venus to the open reading frame (orange boxes) of the Nkx6-1 gene. The loxP-flanked PGK driven neomycin (neo) selection cassette was removed by Cre-recombinase mediated excision. Nkx6-1 5’- and 3’-untranslated regions (UTRs) are indicated by black boxes, the predicted promoter region (yellow box) and transcriptional start sites (TSR, red boxes) are indicated. Primers used for PCR genotyping are designated EP1429, EP1463, EP1499, EP1622 and EP1704. The positions of the homology regions to generate the targeting construct are indicated by dashed lines. (**b, c**) PCR genotyping of Nkx6-1-VFNeo/+ mice using primers 1429, EP1463, EP1622 for the 5’ PCR confirmation of the targeted allele Nkx6-1-VFNeo (545 bp) versus the WT allele (877bp) and the primers EP1463, EP1499 and EP1622 for the 3’ PCR confirmation of the targeted allele Nkx6-1-VFNeo (591 bp) versus the WT allele (877bp). (**d**) PCR primers EP1499, 1622 and 1704 were used to distinguish the allele before (Nkx6-1-VFNeo; 591 bp) and after removal of the neo selection cassette (Nkx6-1-VF; 741 bp). (**e**) Primers EP1463, EP1622 and EP1704 were used to distinguish WT from heterozygous or homozygous mice of Nkx6-1-VF resulting in 877bp for the WT allele and 741 bp for the targeted allele. (**f**) Western blot analysis on lysates from islets of Langerhans using Nkx6.1 antibody to detect the WT protein as double band at approximately 50 and 53 kDa and Nkx6-1-VF protein at 77 and 80 kDa. (**g, h**) Both Flag and GFP antibodies detected the Nkx6-1-Venus Fusion protein at 77 and 80 kDa as well as several degradation products. β-tubulin was used for loading control.

### Spatio-temporal expression pattern of Nkx6-1-VF protein during embryonic pancreas development

To assess whether the Nkx6-1-VF protein reflects the expression pattern of the endogenous Nkx6-1 protein, we stained embryos and embryonic pancreatic sections from the reporter mice. At E9.5, Venus presence was found in pancreatic buds marked by high expression of Pdx1 and Foxa2 (Fig. 2a; yellow arrows) as well as in the neural tube (Fig. 2a; white arrows) as previously reported (Li et al., 2016). At 11.5, Nkx6-1-VF is co-expressed with the TF Pdx1 in the pancreatic epithelium. Interestingly Nkx6-1-VF was found in both pancreatic buds (Fig. 2c) whereas it was absent in the first appearing glucagon secreting endocrine cells during the primary transition (Fig. 2b). Embryonic pancreatic sections of E12.5 embryos presented the fusion protein localized in the cell nucleus and restricted to the trunk domain of epithelium (Fig. 2d; yellow arrow). At E12.5 the Nkx6-1-VF reporter showed low or absent expression in the tip domains of the pancreatic epithelium but was highly expressed in the trunk domains where it colocalized with Pdx-1. Correctly patterned tip and trunk domains indicate that the Nkx6-1-VF does not impair transcription factor function and allows correct patterning of the pancreatic epithelium. The presence of tip and trunk domains indicates proper function of the Nkx6-1-VF TF function in the patterning of the pancreatic epithelium. (Fig. 2d; white arrows). The co-expression of Nkx6-1-VF and Pdx1 was also observed at E16.5 and E18.5. At these stages, we found low expression of the fusion protein in epithelial cells defined by the expression of E-cadherin (Fig. 2e, f; purple arrows). Additionally, Nkx6-1-VF was expressed at higher levels in certain cell populations in close proximity to the ductal epithelium that resembled proto-islets (Fig. 2e; white arrows). Overall, these results demonstrate that the Nkx6-1-VF protein mirrors the endogenous Nkx6-1 spatio-temporal expression pattern during pancreas development.

**Figure 2.**
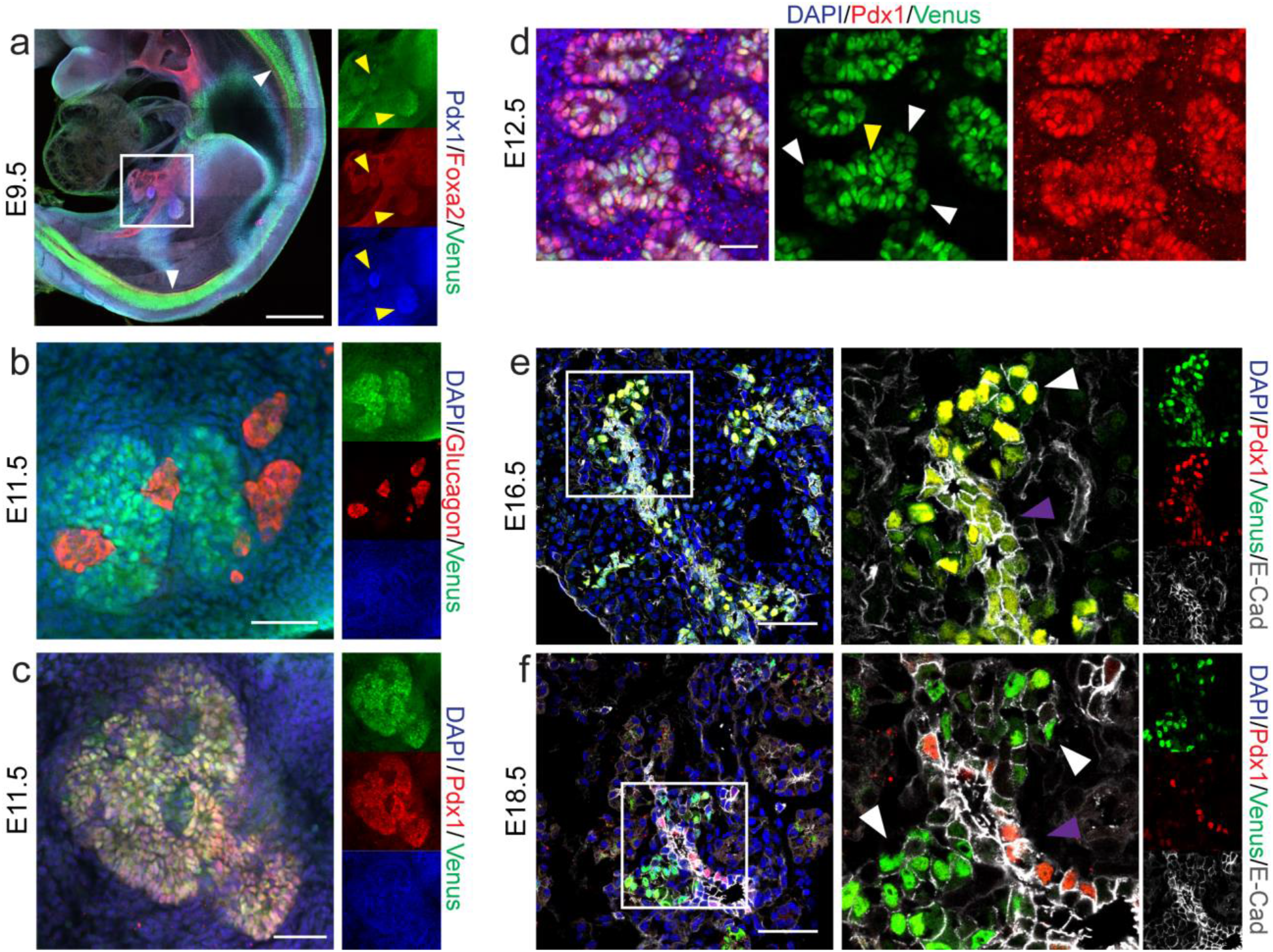
Nkx6-1-VF expression during embryonic pancreas development. (**a**) Whole embryo immunostaining using iDSCO protocol of E9.5 presenting Nkx6-1-VF expression in notochord (white arrows) and pancreatic buds marked by high expression of Pdx1 and Foxa2 (yellow arrows). Tile-Scan; Size bar 500 μm. (**b, c**) Whole pancreas immunostaining at stage E11.5 shows (**c**) colocalization of Nkx6-1-VF and Pdx1 in the pancreatic epithelial but (**b**) absent in the glucagon-secreting cells. Scale bar 50 μm. (**d, e, f**) Pancreas section immunostaining of E12.5, E16.5 and E18.5 analyzing the expression of Nkx6-1-VF during embryonic pancreas development. The expression of the fusion protein through primary and secondary transition follows a similar pattern as Pdx1. (**d**) At E12.5 the expression of Nkx6-1-VF is observed in the duct domain (yellow arrow) but not in the tip domain (white arrow). Scale bar 20 μm. (**e, f**): During endocrine lineage specification higher expression levels outside the duct marked with high expression of membrane marker E-cadherin is observed. Scale bar 50 μm.

### Nkx6-1-VF marks the endocrine lineage during secondary transition

During the secondary transition of pancreas development the expression of Nkx6-1 is found in the ductal epithelium and endocrine progenitor cells. However, the levels of this TF increase and become restricted to β-cells (Øster et al., 1998). Therefore, we evaluated the expression pattern of Nkx6-1-VF in pancreatic sections at E16.5 and E18.5, when endocrinogenesis occurs. Using immunohistochemical analysis, we found low expression levels of Nkx6-1-VF in Sox9^+^ ductal cells and increased expression of the fusion protein in a Sox9^−^ cell clusters close to the trunk epithelium (Fig 3a-c). To define the identity of the later population, we performed co-staining of the Nkx6-1-VF with insulin and glucagon to identify α- and β-cells in E18.5 pancreatic sections. The results indicated expression of Nkx6-1-VF in β-cells but absent of very low residual levels in α-cells (Fig 3d, e). Furthermore, co-staining of the fusion protein with the exocrine marker α-amylase revealed no expression of the Nkx6-1-VF protein in the exocrine compartment (Fig 3f). Collectively, these data confirm the expression of Nkx6-1-VF protein in the ductal bipotent progenitors and then specific to β-cells during secondary transition of pancreas development.

**Figure 3.**
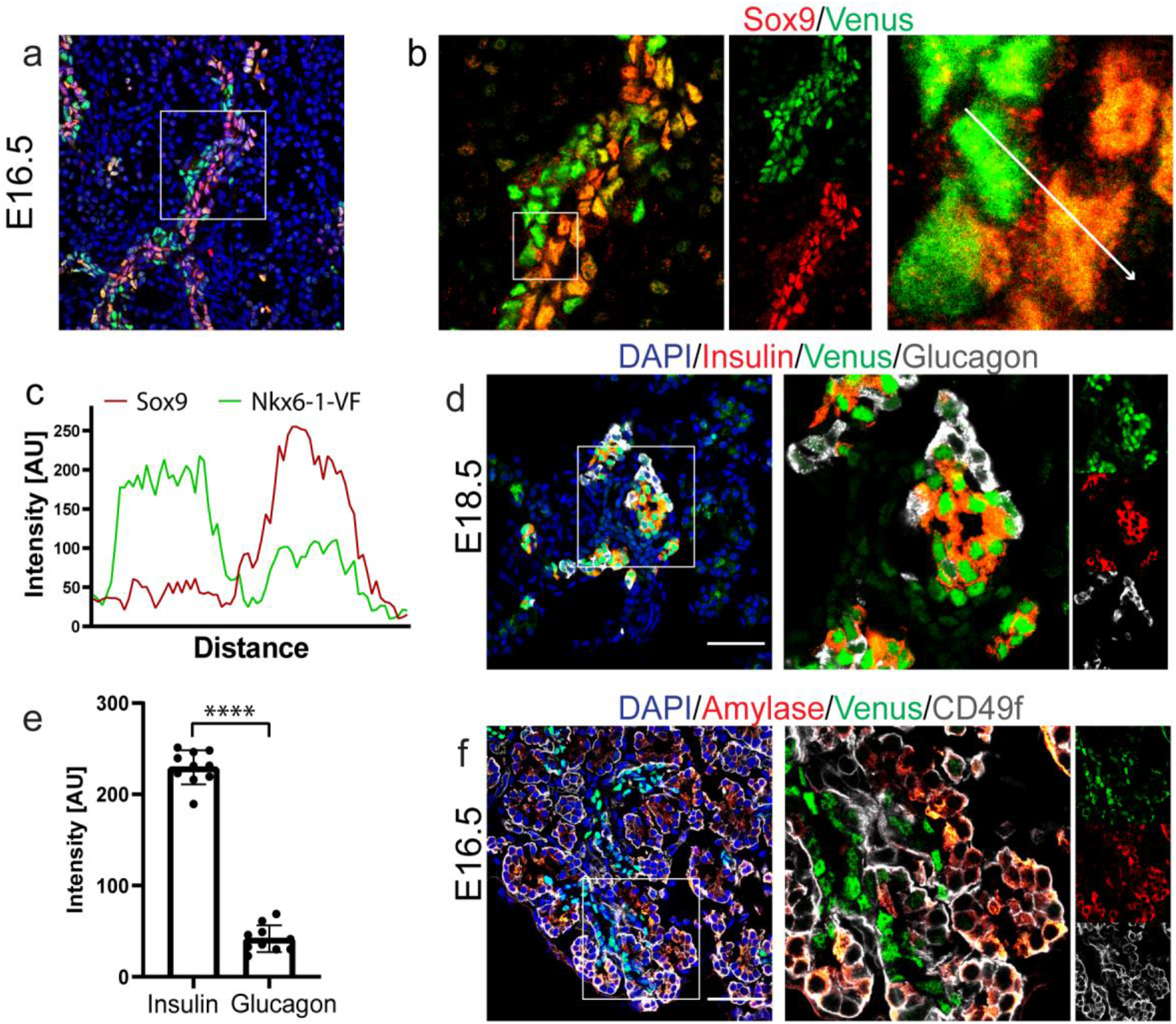
Nkx6-1-VF marks the endocrine lineage during secondary transition. (**a, b, f**) Immunostaining of E16.5 and (**d**) E18.5 pancreas section exhibiting Nkx6-1-VF expression during endocrine cell formation and β-cell lineage specification. (**b**) Immunostaining analysis of E16.5 pancreas shows that Nkx6-1-VF expression is express in low levels in the duct (Sox9^+^/Nkx6-1-VF low) and high levels near the duct (Sox9^−^/ Nkx6-1-VF high), (**f**) but not in the exocrine cells marked by amylase (**d, e**). Nkx6-1-VF high expression levels correlate with endocrine lineage formation marking specifically insulin secreting cells as observed at E18.5. Scale bar 50 μm

### Nkx6-1-VF expression pattern in the adult pancreas

In adult islets, Nkx6-1 is exclusively expressed in β-cells and not in any other endocrine cell types (Jensen et al., 1996). To investigate whether the Nkx6-1-VF reporter remains active in the islet of Langerhans, we performed immunostaining of sections derived from adult pancreas or isolated islets. We co-stained the sections with antibodies against Venus, Insulin and Nkx6-1. Venus expression was highly co-localizing with Nkx6-1 (Fig. 4a, f), but also low Venus levels were found within the cytoplasm of exclusively β-cells (Fig. 4a). This was further confirmed by staining of Venus with Glucagon, indicating the presence of no Venus nuclear signal in α-cells (Fig. 4b). Weak cytoplasmatic localization of Venus reporter in β-cells is likely caused by protein degradation and the Venus fragments still being immunoreactive (Genové et al., 2005). In addition, we analyzed the expression of the β-cell maturation marker, Urocortin 3 (Ucn3) in the Nkx6-1-VF-expressing cells. At postnatal day 3 (P3), when the majority of β-cells are still immature, we found the expression of Ucn3 only in a fraction of Nkx6-1-VF-expressing cells (Fig. 4c). On the contrary, at P45 all the Nkx6-1-VF^+^ cells expressed Ucn3 (Fig. 4d), indicating that the fusion reporter protein does not hamper β-cell maturation. Next, we performed time-lapse imaging of the isolated islets derived from Nkx6-1-VF mice. Fluorescent intensity of the Nkx6-1 reporter was sufficient to track single β-cells during time-lapse imaging and follow the β-cell movement (Fig. 4e; Suppl. movie). Additionally, as the Nkx6-1-VF mRNA uses the endogenous Nkx6-1 UTR, it may be used as a sensor to study miRNA function. Finally, we successfully sorted Nkx6-1-VF^+^ cells from the isolated adult islets using FACS (Fig. 4g), indicating the capability of the fusion reporter protein for specific isolation of β-cells.

**Figure 4.**
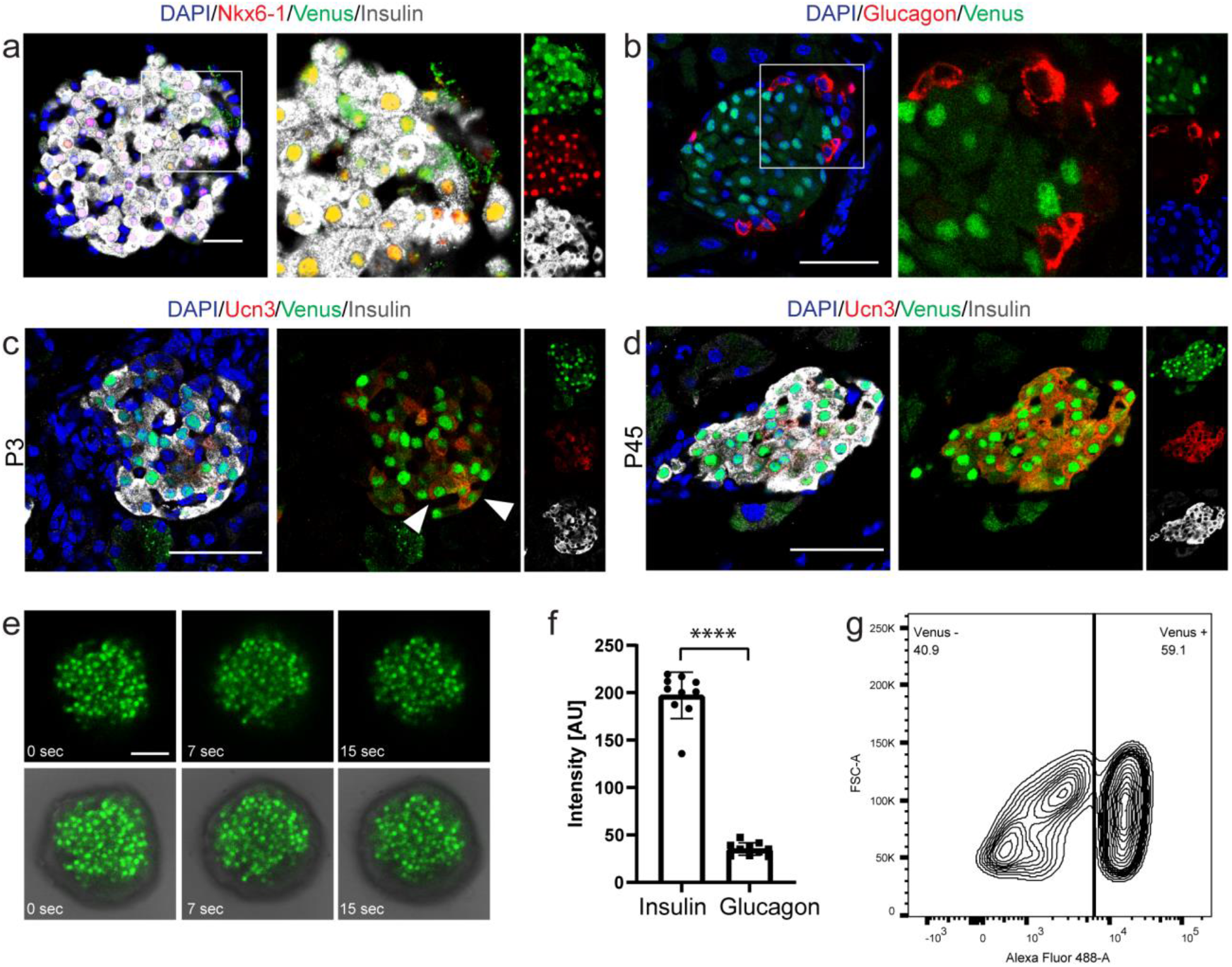
Nkx6-1-VF adult mice express Venus in the mature islets and can be used to sort β-cells *in vivo*. (**a**) Nkx6-1-VF correlates with Nkx6-1 and insulin-producing cells (**b, f**) but not with glucagon-secreting cells. (**c, d**) Immunostaining of P3 and P45 pancreatic sections showing the maturation of Nkx6-1-VF β-cells. (**c**) At P3 only a fraction (white arrowheads) of the reporter cells express Ucn3, (**d**) while all the reporter cells express this maturation marker at P45. Scale bar 50 μm. (**e**) Time-lapse imaging of isolated islets from adult Nkx6-1-VF mice. Scale bar 50 μm. (**g**) Representative FACS plots indicating the successful separation of endocrine cells from the isolated adult islets based on the Venus fluorescent signal.

In summary, we have generated the first Nkx6-1-Venus Fusion reporter mouse model that resembles the expression of endogenous Nkx6-1 and provides a unique tool for isolation of Nkx6-1-expressing pancreatic cells at different stages. Therefore, this mouse line offers a valuable technical support to study pancreas development and β-cell function in health and disease.

## Material and methods

### Generation of the targeting construct

For the targeting vector, 5’ homology region (HR) and 3’ HR were PCR amplified using C57B16 BAC (RPCIB-731L18311Q) as template and using primers as follows: EP_1197 and EP_1198 primers (see table1) for 5’ HR and EP_1199 and EP_1200 primers for 3 ́ HR. 5’ HR was subcloned via NotI and XbaI and 3’ HR was subcloned via HindIII and XhoI, into the pBluescript KS (pBKS), generating the pBKS-Nkx6-1 Ex3-HR. Using primers EP_1126 and EP_1201 on a Venus containing DNA template a Venus-3xFlag tag fragment (819 bp) was amplified and gel purified after XbaI and SpeI digestion and subcloned between 5’ and 3’ HRs, resulting in pBKS-Nkx6-1 Ex3-HR-Venus-3xFlag. The PGK promoter-driven neomycin resistance gene flanked by loxP sites (loxP-Neo-loxP) was released by BamHI and EcoRI digestion from the PL452-loxP (Copeland et al., 2001) and cloned into these sites downstream of the Venus gene resulting in the targeting vector pBKS-Nkx6-1 Ex3-HR-Venus3xFlag-Neo. Two gRNA sequences targeting up- and downstream near the stop codon of Nkx6-1 (Fig1a) were selected using online Crispr resources (Hsu et al., 2013). To generate CRISPR expression vectors self-annealed oligos (Nkx6-1 Crispr #11 and #16 fwd and rev; Table1) duplex with BbsI overhangs were cloned into BbsI digested pBS-U6-chimericRNA (a generous gift from O. Ortiz, Institute of Developmental Genetics, Helmholtz Zentrum München) resulting in pBS-U6-chimericRNA Nkx6-1#11 and #16. Successful integration of CRISPRs into pbs-U6-chimericRNA vectors was confirmed by sequencing.

**Table 1.**
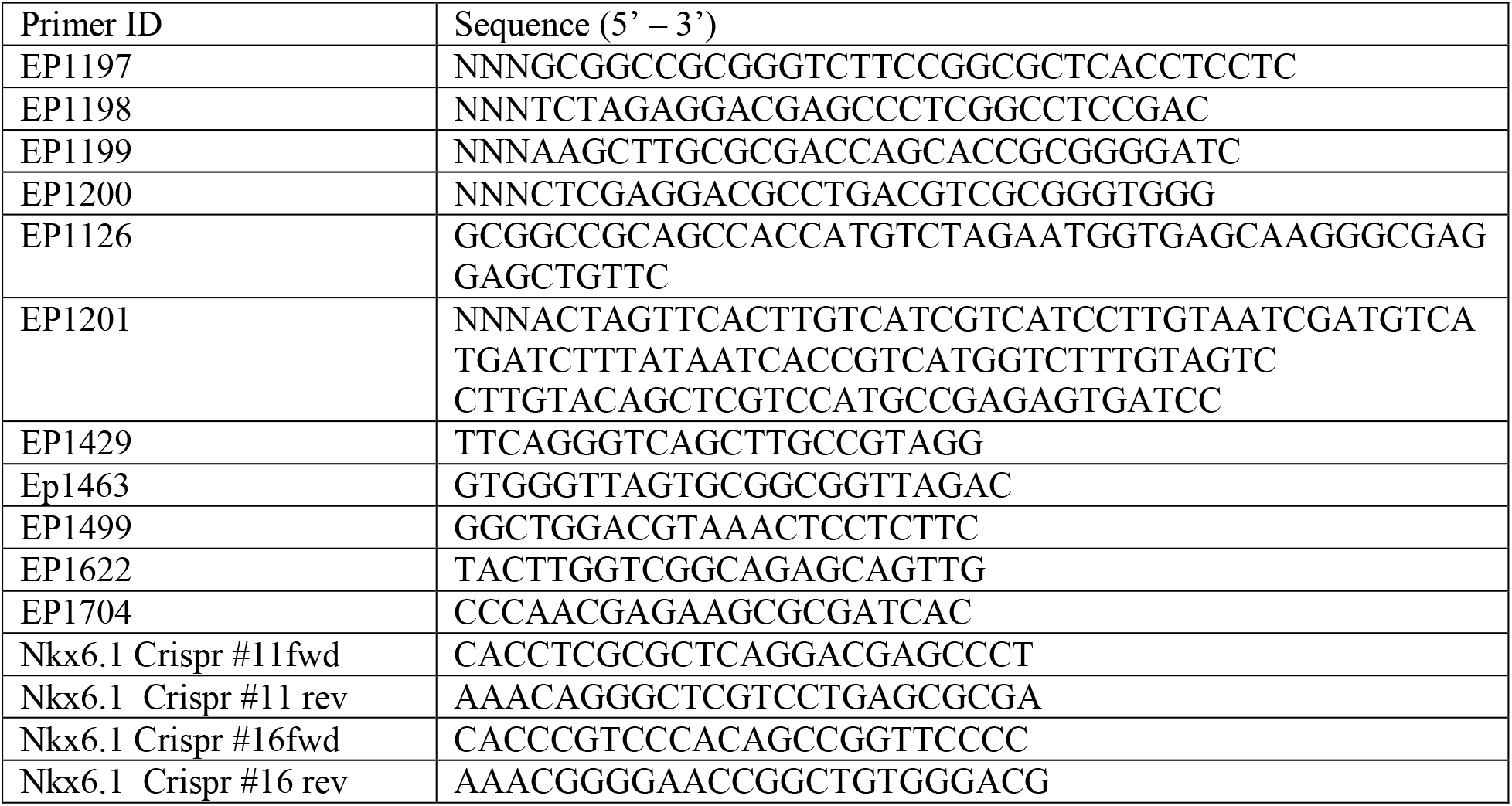
Primers sequences

### Cell culture and homologous recombination in ES cells and mouse generation

Mouse ES cells were cultured on a murine embryonic feeder (MEF) layer in Dulbecco’s Modified Eagle Medium (DMEM, Invitrogen, Carlsbad, CA) containing 15% fetal calf serum (FCS, PAN, Aidenbach, Germany), 2 mM L-glutamine (Invitrogen, 200 mM), non-essential amino acids (Invitrogen, Carlsbad, CA, 1003), 100 μM b-mercaptoethanol (Invitrogen, 50 mM), and 1500 U/ml leukaemia inhibitory factor (LIF, Chemicon-Millipore, Billerica, MA, 107 U/ml). Cells were split every two days using trypsin (0.05% trypsin, 0.53 mM EDTA; Invitrogen). IDG 3.2 ES cells (Hitz et al., 2007) were electroporated with a mixture of pBKS-Nkx6-1 Ex3-HR-Venus3xFlag-Neo targeting vector, both pBS-U6-chimericRNA Nkx6-1#11 and #16 as well as Cas9 nickase overexpression vector (pCAG Cas9v2D10A-bpA; a generous gift from O. Ortiz, Institute of Developmental Genetics, Helmholtz Zentrum München). Neo resistant clones were selected using 300 μg/ml G418 (Invitrogen, 50 mg/ml). Homologous recombination at the Nkx6-1 locus was confirmed by homology arm spanning PCRs. Homologous recombined ES cell clones were aggregated with CD1 morulae and the resulting chimeras gave germline transmission of the Nkx6-1VFneo allele. The floxed neo selection marker cassette was deleted in the germ line by intercrossing with the ROSA-Cre mouse line.

### Genotyping

Mice genotyping was assessed by PCR analysis of ear clip-derived DNA as template. The excision of the Neo cassette was confirmed through the genotyping of PCR using the primers EP 1499, 1622 and 1704 generating a 591 bp product for the Nkx6-1^Venus Neo^ allele and a 741 bp product for the ^Venus delta Neo^ allele (Fig. 1b). To genotype the homozygous and heterozygous Nkx6-1-VF mice, PCR analysis was performed at 60 °C annealing temperature using the primers EP 1463, 1622 and 1704. The WT mice (+/+) generated a 877 bp PCR product, distinguished from the 741 bp band of the homozygous mice (V/V). Two products of 877 and 741 bp were identified for the heterozygous mice (V/+) (Fig. 1b).

### Western blot analysis

Western blot analysis was performed according to the standard protocols. Briefly, lysates from pancreatic Islets of Langerhans were subjected to the SDS-PAGE electrophoresis and transferred to the nitrocellulose membranes. After blocking, the membranes were incubated with anti-GFP (Rabbit 1:2000; Invitrogen A11122), anti-Nkx6-1 (Goat 1:300; R&D Systems AF5857), anti-Flag (Mouse HRP, Sigma A8592; 1:10.000) and anti–GAPDH (Mouse 1 μg/ml; Merck/Millipore). HRP-conjugated secondary antibodies were used as follows: Anti-Mouse HRP (1:10000; Millipore; 12-349), anti-Rabbit HRP (1:10.000; Dianova, 111-035-046) and Anti-Goat HRP (1:10.000; Dianova 305-035-045). The signals were detected by enhanced chemiluminescence (Thermo Scientific).

### Pancreas dissection

Embryonic or adult pancreata were dissected and fixed in 4% PFA in PBS 2 hrs at RT. The tissues were then cryoprotected in 10% and 30% sucrose solutions for 2 hrs at RT and finally incubated in 30% sucrose and tissue embedding medium (Leica) (1:1) at 4 °C overnight. Afterwards they were embedded in a tissue-freezing medium (Leica) and stored at −80 °C. Sections of 20 μm thickness were cut from each sample, mounted on a glass slide (Thermo Fisher Scientific) and dried for 10 min at room temperature before use or storage at −20 °C.

### Immunostaining of sections

Cryosections were rehydrated with 1x PBS and permeabilized with 0.2% Triton X-100 in 0.1M Glycine solution for 30 min. Samples were then incubated in a blocking solution (10% PCS, 3% Donkey serum, 0.1% BSA and 0.1% Tween-20 in PBS) for 1 hr at RT. Afterwards primary antibodies diluted in blocking solution were added to the samples overnight at 4 °C. Following primary antibodies were used for staining: anti-ucn3 (Rabbit 1:300; Phoenix Pharmaceuticals H-019-29), anti-glucagon (Guinea pig 1:2500; TAKARA M182), anti-GFP (Chicken 1:1000; Aves Lab; GFP-1020), anti-insulin (Rabbit 1:300, Cell signaling 3014), goat anti-Pdx1 (Goat 1:300; Abcam AB47383), anti-E-Cadherin (Rat 1:300; Kremmer SC-59778), anti-Cd49f (Rat 1:300; BD 555734), anti-Sox9 (Rabbit 1:300; Abcam AB5535), anti-Amylase (Rabbit 1:300; Abcam AB21156), anti-Nkx6-1 (Goat 1:200; R&D Systems AF5857). Primary antibodies were washed with 1x PBS and secondary antibodies diluted in blocking solution were added 4 hrs at RT: anti-Rabbit 555 (1:800; Invitrogen; A31572), anti-Chicken Cy2 (1:800; Dianova, 703-225-155), anti-Guinea Pig 649 (1:800; Dianova; 706-495-148), anti-Rat DyLight 549 (1:800; Dianova, 712-505-153) and anti-Goat 555 (1:800; Invitrogen A21432). Images were obtained with a Leica microscope of the type DMI 6000 using the LAS AF software.

### Islet isolation

Islet isolation was performed by digestion of adult pancreas as described previously (Bastidas-Ponce et al., 2019). Collagenase P (Sigma-Aldrich, Germany) dissolved in Hanks Balanced Salt Solution (HBSS) with Ca2^+^/Mg2^+^ was injected into the bile duct to perfuse the pancreas After a gradient preparation (5 mL 10% RPM + 3 mL 40% Optiprep/ per sample), islets were handpicked and incubated at 37 °C 5% CO2 in culture with 11 mM glucose in RPMI medium 1640 supplemented with 10% (vol/vol) FBS Heat Inactivated, 1% (vol/vol) penicillin and streptomycin.

### FACS analysis

Islets from adult Nkx6-1-VF homozygous mice were disaggregated into single cells with TriplE 10 min at 37 °C and resuspended in FACS buffer (PBS with FCS 10%) and filtered through a 35 mm cell strainer. Cells were analyzed and isolated using an Aria III (BD Biosciences).

### Time-Lapse Live Imaging

Time-lapse imaging was carried out as described by Burtscher & Lickert, 2009. Islets were incubated in RPMI medium 1640 supplemented with 10% (vol/vol) FBS Heat Inactivated, 1% (vol/vol) penicillin and streptomycin cultured on glass-bottom dishes in a 37 °C incubator with 5% CO2. To avoid evaporation, the medium was covered with mineral oil. Image acquisition was performed on a Leica DMI 6000 confocal microscope equipped with an incubation system and image analysis was carried out using Leica LAS AF software.

### iDisco for clearing of mouse embryos

E9.5 mouse embryos were processed according to the iDisco protocol published (Renier et al., 2014). Primary antibodies were incubated for 5 days, secondary antibodies for 4 days at 37°C. Following primary antibodies were used for stainings: anti-GFP (Goat 1:500, Biotrend; 600-101-215), anti-Foxa2 (Mouse 1:500; Millipore 17-10258), anti Pdx1 (Rabbit 1:500; Cell Signaling; D59H3) and secondary antibodies: anti-Goat 488 (1:800; Invitrogen A11055), anti-Mouse CY5 (1:800; Dianova; 715-175-151) and anti-Rabbit 555 (1:800; Invitrogen A31572). Images were taken using tile scan mode on a Zeiss LSM 880 using ZenBlack software.

## Supporting information

Supplementary movie 1 GFP and BF

Supplementary movie 1 GFP

## Acknowledgment

We thank Jessica Jaki and Aimée Bastidas-Ponce for their technical support. This work was supported by the Helmholtz-Gemeinschaft (Helmholtz Portfolio Theme ‘Metabolic Dysfunction and Common Disease) and Deutsches Zentrum für Diabetesforschung (DZD).

